# 3D human skeletal muscle organoids reveal distinct effects of high-dose dihydronicotinamide riboside on muscle development

**DOI:** 10.64898/2025.11.28.690663

**Authors:** Ashwin Venkateshvaran, Swarang Sachin Pundlik, Yavanica Suresh, Akshay Hegde, Bhushan Venkatesh, Srujanika Rajalakshmi, Sikhasmita S Dowari, Alok Barik, Heera Lal, SriCharan Lakkaraju, Nikhil Zakariah, Gautham Pranesh, G Mani Subramanian, Arvind Ramanathan

## Abstract

The evaluation of NAD+-boosting compounds in human skeletal muscle is hindered by limitations of traditional 2D cultures and animal models. Human-relevant, three-dimensional (3D) engineered skeletal muscle organoids offer a promising platform to assess the biological effects of metabolic modulation. Here we engineered 3D human skeletal muscle organoids to investigate the impact of dihydronicotinamide riboside (NRH), a potent NAD+ precursor. Sustained exposure to high NRH concentrations (500 µM) enhanced early differentiation markers, including increased myotube fusion and fast-twitch fiber area, but concurrently induced structural defects such as disrupted sarcomeric organization, enlarged acetylcholine receptor clusters, and impaired acetylcholine-stimulated calcium signaling. These results reveal that excessive and sustained NAD+ elevation can uncouple rapid differentiation from proper maturation in muscle tissue. Our results highlight the importance of dose and duration optimization for NAD+-boosting compounds and establish 3D engineered muscle organoids as a valuable non-animal platform for mechanistic toxicology and preclinical safety assessment.

## Introduction

There is a growing need for human-relevant, non-animal platforms capable of detecting toxicological responses to nutraceuticals, pharmaceuticals, and metabolic modulators. Classical animal studies are expensive, slow, and often limited in their ability to predict human-specific muscle toxicity. 2D cell cultures lack the architectural and functional sophistication required to capture tissue-level maturation. As regulatory agencies increasingly encourage New Approach Methodologies (NAMs) (Sewell, Alexander-White et al. 2024, Ouedraogo, Alepee et al. 2025), engineered 3D organoid and micro physiological systems have emerged as powerful pre-clinical platforms. They bridge pre-clinical assays and whole-animal toxicology (Bridgeman, Pamies et al. 2025). Skeletal muscle is particularly underserved in this context despite being a major target for metabolic and longevity-associated compounds (Demontis, Piccirillo et al. 2013, Sharples, Hughes et al. 2015, Goody and Henry 2018, Morgan and Partridge 2020). Muscle function depends on aligned multinucleated fibers, mature sarcomeres, acetylcholine receptor (AChR) clustering, mitochondrial organization, and calcium handling. These features do not arise in 2D monolayers but develop in 3D engineered constructs. These properties enable direct measurement of structural integrity and functional performance, providing a physiologically human-relevant platform for toxicology assessment.

Recent advances in muscle micro physiological systems (MPS) and organ-on-chip devices have demonstrated the feasibility of generating contractile human muscle tissues capable of force measurement, electrical stimulation, and drug response profiling (Bian, Juhas et al. 2012, Madden, Juhas et al. 2015, Gholobova, Gerard et al. 2018, Khodabukus, Madden et al. 2019, Parafati, Giza et al. 2023). Platforms such as soft-post tissues (Franken, van der Wal et al. 2025), perfused microfluidic chips (Parafati, Giza et al. 2023), and neuromuscular co-cultures (Afshar Bakooshli, Lippmann et al. 2019) have been used to model muscular dystrophies and screen bioactive molecules (Tejedera-Villafranca, Montolio et al. 2023). However, comparatively few studies have applied these systems to metabolic toxicology, and even fewer have examined how extreme metabolic challenges affect coordinated sarcomere organization and fiber-type composition. Our work builds on and complements these efforts by demonstrating that 3D engineered human muscle (ESM) is highly sensitive to metabolic and intracellular redox perturbations and can detect functional deficits that are not clearly measurable in 2D culture. This is relevant for NAD+-boosting molecules, which are widely explored for their effects on metabolism and aging (Verdin 2015, Fang, Lautrup et al. 2017, Katsyuba and Auwerx 2017, Rajman, Chwalek et al. 2018). Dihydronicotinamide riboside (NRH) is a potent precursor, capable of elevating NAD+ levels more rapidly and several fold higher than nicotinamide riboside (NR) or nicotinamide mononucleotide (NMN) (Giroud-Gerbetant, Joffraud et al. 2019, Yang, Mohammed et al. 2019). While these properties make NRH of interest as a metabolic intervention, the consequences of rapid, substantial and sustained NAD+ elevation on human skeletal muscle remain poorly understood. This is particularly relevant in the context of tissue-level processes such as sarcomere assembly, calcium signaling, and fiber-type specification, which are governed by strict energetic and structural requirements.

In this study, we use ESMs to evaluate the upper-bound effects of sustained NRH exposure. By integrating morphological, fiber-type, mitochondrial, and calcium-based functional endpoints, we demonstrate that this non-animal system can simultaneously detect both early differentiation changes and later-stage structural and functional alterations, including those related to calcium handling. These findings position ESMs as a valuable New Approach Methodology (NAM) for mechanistic toxicology and highlight its potential role in evaluating both dose and exposure response relationships of metabolic compounds, nutraceuticals, and bioactive small molecules in a tissue-specific context.

## Materials and methods

### Fabrication of PDMS scaffolds

3D scaffolds for ESM culture were designed in AutoCAD. The design was exported as an STL file and used to manufacture polyurethane (PU) masters. PU blocks were laser-cut to generate the master mould (positive) and then cleaned thoroughly with 70% ethanol to remove particulates and residual debris. The PU masters were plasma-treated in an O□plasma cleaner (Harrick) for 5 min to activate the surface. The activated surface was then functionalised with vapour-phase deposition of Trichloro(1H,1H,2H,2H-perfluorooctyl) silane in a commercially acquired vacuum desiccator at 150 mbar to reduce adhesion and facilitate release from the PU mould.

Ecoflex negatives were again plasma-treated and silanized. Sylgard 184 polydimethylsiloxane (PDMS, Dow Corning) precursor and curing agent were mixed at a 10:1 ratio (w/w), degassed under vacuum, and poured into the Ecoflex moulds. This was left overnight, and 15:1(Base: curing agent) polydimethylsiloxane (Sylgard 154) was poured into the negatives and left to cure at room temperature for 4 days. The PDMS positives were then manually peeled, cleaned with 70% ethanol, autoclaved, and stored. Each PDMS scaffold contained four rectangular wells, each with a pair of cylindrical posts (0.4 mm diameter × 3.76 mm height) separated by 4.68 mm centre-to-centre. To prevent slippage of the engineered muscle tissues during culture, posts were capped with custom 3D-printed ABS caps. The caps were fixed using small amounts of uncured Ecoflex, cured at 90 °C, autoclaved for sterilization, and stored in sterile Milli-Q water until use.

### Human primary myoblast culture

Human skeletal muscle myoblasts were obtained from Lonza (21-year-old male donor) and Cook MyoSite (29-year-old male donor) and were maintained in DMEM skeletal muscle growth medium (SKMGR) at 37 °C and 5% CO□. For experiments, the cells were trypsinized with 0.125% trypsin–EDTA (Gibco) and seeded at the required densities in cell culture dishes. All cultures tested negative for mycoplasma using the Mycoalert Mycoplasma Detection Kit (Lonza). Passages between 4 and 6 were used.

### Organoid fabrication

P4 myoblasts were trypsinized, counted, and stored on ice after cell counting. A total of 4 × 10□myoblasts were mixed with hydrogel component A (20 µL SKMGR + 2 µL 50 U/mL thrombin). Hydrogel component B contained 10 µL SKMGR + 10 µL 20 mg/mL human fibrinogen + 5 µL Geltrex. Components A and B were kept separately on ice. Component B was mixed thoroughly and added to component A, which was mixed again gently.

A total of 47 µL of the hydrogel–cell mixture was then deposited into each well and allowed to polymerize at 37 °C for 60 min, after which 5 mL of SKMGR containing 1.5 mg/mL aminocaproic acid was added and organoids were allowed to proliferate for 4 days. After 4 days of proliferation, skeletal muscle differentiation medium with 2 mg/mL aminocaproic acid was added, and organoids were allowed to differentiate for 14–16 days with medium changes every alternate day. Bright-field images were acquired at defined time points (P-1, D0, D2, D4, D8, D12, D14).

### NRH preparation and storage

A sterile solution of NRH was prepared at a concentration of 50mg/ml in a final volume of 25ml. A dry volume of 1250mg of NRH was dissolved in a Na2hPO4 buffer and the final pH was maintained at 9.0. The solution was stored at °80degrees before use.

### NRH treatment of engineered skeletal muscle organoids

ESMs after the proliferation stage were washed three times with DPBS. The appropriate amount of NRH in vehicle (Sodium Biphosphate Buffer) was added to skeletal muscle differentiation medium and applied to the ESMs. Medium containing NRH was replenished every 48 h, and organoids were harvested at the indicated time points.

### Fixation, cryoprotection, and sectioning

Organoids were rinsed in DPBS and fixed in 4% paraformaldehyde (PFA) overnight at 4 °C, washed 3× with PBS, cryoprotected in 30% sucrose overnight at 4 °C, embedded in OCT, frozen on dry ice, and cryosectioned (Leica CM 1860 Cryotome) at 10–12 µm. Sections were mounted on standard microscopy slides (Blue Star). Slides were air-dried, outlined with a hydrophobic barrier pen, and stored at –80 °C before immunostaining.

### Immunostaining

Organoid cryosections were equilibrated in room temperature for 10–15 min and fixed in 4% PFA for 10 min. Sections were washed twice with 1× PBS for 5 min each. Permeabilization was performed using 0.4% Triton X-100 in PBS (PBS-T), with incubations of 3 washes of 10 min at room temperature. Sections were then blocked using 5% Fetal bovine serum (FBS) prepared in PBS-T for 30 min at room temperature.

Following blocking, sections were incubated with primary antibodies diluted in 1% serum in PBS for 1 h at room temperature. After incubation, sections were washed twice with 1× PBS-T for 10 min each. Secondary antibodies diluted in 1% serum in PBS were then applied for 1–2 h at room temperature, followed by two additional 10-min washes in 1× PBS-T. Nuclear counterstaining was performed using DAPI (1:1000) for 10 min. After a final wash, sections were mounted with antifade mounting medium (Prolong Gold, Invitrogen) and imaged using fluorescence microscopy.

### Microscopy

Widefield epifluorescence imaging was performed on a Nikon Eclipse inverted fluorescence microscope (Nikon, Tokyo, Japan) with LED illumination and 10×, 20×, and 40× Plan Fluor objectives (0.3–0.75 NA. Confocal imaging and higher-magnification imaging were performed on an FV3000 laser-scanning confocal microscope (Olympus, Tokyo, Japan) with appropriate excitation/emission filter sets for DAPI, FITC/Alexa 488, TRITC/Alexa 568, and far-red fluorophores. For each experiment, exposure times, laser powers, and detector gain were kept constant across treatment groups.

### Quantitative image analysis

All image analysis was performed using FIJI/ImageJ (version 1.54) with the MorphoLibJ and OrientationJ plugin suites and custom macros.

### Myotube alignment, nuclear area, and nuclear shape

For longitudinal sections stained with pan-MYHC and DAPI, channels were thresholded (Otsu, Huang, or manually adjusted) to generate binary masks. Segmentation and labelling of nuclear metrics were performed using MorphoLibJ. OrientationJ’s structure tensor analysis was applied to the MYHC channel to calculate the local orientation of myofibres and generate orientation histograms. Myotube thickness was calculated using the Myotube Analyser.

Nuclear area and nuclear shape descriptors (convexity, perimeter, circularity) were quantified from the DAPI mask using MorphoLibJ’s “Shape Features” module. For each ESM, three entire longitudinal sections were analysed; three sections per ESM were used.

### Fusion index

Fusion index was quantified on longitudinal sections stained for pan-MYHC, DAPI, and AChR using the MATLAB application “Myotube Analyser,” which integrates MYHC and DAPI masks to determine the number of nuclei within MYHC-positive myotubes. Briefly, MYHC-positive regions were segmented and dilated slightly to create myotube masks, which were overlaid on the DAPI channel. Nuclei whose seed watershed masks fell within the myotube mask were classified as fusion positive. Fusion index was calculated as the percentage of total nuclei per FOV that were fused. At least 30 FOVs per condition (vehicle vs NRH) were analysed across multiple organoids and donors.

### Myotube number, cross-sectional area, and total transverse area

For transverse sections stained with F-actin, the actin channel was smoothed and converted into a binary mask. Individual myotubes were segmented, and three parameters were computed: (1) number of myotubes per section; (2) mean myotube cross-sectional area (µm^2^); and (3) total transverse area occupied by myotubes (sum of all myotube areas). Each data point represented a single FOV; ≥30 FOVs per condition were analysed.

### Fibre-type composition

Transverse sections stained for MYH2, MYH7, F-actin, and DAPI were analysed to quantify fibre-type-specific areas. MYH2- and MYH7-positive regions were segmented using channel-specific thresholds, and the area of each isoform-positive mask as well as its percentage of the total tissue cross-sectional area were computed. A minimum of 10 FOVs per condition were analysed across donors.

### Sarcomere spacing

For sarcomeric α-actinin–stained longitudinal sections, line profiles were drawn along the longitudinal axis of myotubes. Fluorescence intensity profiles were saved as CSV files, and peak-to-peak distances between α-actinin bands were measured using a custom ImageJ macro. At least five sarcomeres per measurement and five measurements per organoid were quantified.

### AChR cluster analysis

Longitudinal sections co-stained for MYHC and AChR (α-bungarotoxin) were used to quantify AChR cluster size. AChR puncta were segmented using size and intensity thresholds designed to exclude background. Individual AChR clusters were labelled, and their areas were measured. Values were averaged per section; each data point represented one full 60× FOV, with ≥30 FOVs per condition analysed.

### C2C12 cell culture

C2C12 murine myoblasts were maintained in high-glucose DMEM supplemented with 10% fetal bovine serum (FBS) and 1% penicillin–streptomycin at 37 °C in a humidified incubator (5% CO□). Cells were passaged at 60–80% confluence using 0.05% trypsin–EDTA. For all assays, cells between passages 5–15 were used. For differentiation, confluent cultures were switched to differentiation medium (DMEM containing 2% horse serum and 1% penicillin–streptomycin) and maintained for 4–6 days with medium changes every 48 h.

### C2C12 viability assay (MTT)

To assess NRH cytotoxicity, C2C12 myoblast viability was quantified using an MTT assay. C2C12 cells were seeded into 96-well plates at 5 × 10^3^–1 × 10□cells per well in growth medium and allowed to attach overnight. When cultures reached ∼80–90% confluence, medium was replaced with differentiation medium containing vehicle or NRH (125, 250, or 500 µM). Treatments were maintained for 24, 48, or 72 h, with one medium change (fresh NRH or vehicle) at 48 h for the 72-h condition.

At the indicated time points, 0.5 mg/mL MTT (3-(4,5-dimethylthiazol-2-yl)-2,5-diphenyltetrazolium bromide) in differentiation medium was added to each well and incubated for 3–4 h at 37 °C. Supernatants were aspirated, and formazan crystals were solubilized in 100– 150 µL DMSO (or isopropanol) per well. Absorbance was measured at 570 nm.

### C2C12 fusion assay

C2C12 myoblasts were seeded on glass coverslips placed in 24-well plates (≈3 × 10□–5 × 10□cells per well) in growth medium and allowed to reach full confluence. Cultures were then switched to differentiation medium with vehicle or NRH (125 or 250 µM) and maintained for 4–5 days, with medium (and NRH) refreshed every 48 h.

Cells were washed with PBS and fixed in 4% paraformaldehyde for 15 min at room temperature. Following PBS washes, cells were permeabilized with 0.1–0.2% Triton X-100 in PBS for 10 min and blocked in 5% FBS in PBS-T for 1 h. Coverslips were incubated overnight at 4 °C with a primary pan–myosin heavy chain antibody, washed, and then incubated with an appropriate fluorophore-conjugated secondary antibody for 1 h at room temperature. Nuclei were counterstained with DAPI (1:1000) for 5–10 min. Coverslips were mounted using Prolong Gold antifade mounting medium and imaged by confocal microscopy.

Fusion index was calculated as follows: myotubes were defined as MYHC-positive cells containing ≥2 nuclei, and fusion index was expressed as the percentage of total nuclei per field that resided within MYHC-positive multinucleated myotubes. For each condition, three 12-mm coverslips per experiment were analysed.

### Statistical analysis

All data are presented as mean ± standard deviation (SD) unless otherwise specified. For comparisons between two groups (vehicle vs NRH), two-tailed unpaired Student’s t-tests were used. For multi-group comparisons (e.g., nuclear shape across organoids O1–O4), one-way ANOVA followed by appropriate post hoc tests was applied. Significance thresholds were set at p < 0.05. Exact p-values and the number of replicates for each analysis are reported in the figure legends. Graphs and statistical analyses were generated using GraphPad Prism 9.53.

## Results

### Design and Fabrication of PDMS Scaffolds

The 3D scaffolds were designed using AutoCAD software. The .stl file was then used to laser-cut solid polyurethane (PU) blocks. The PU masters were cleaned with isopropanol to. This formed the primary master, which was then used to cast the Ecoflex (00-20 Shore hardness) negative molds. Ecoflex was chosen due to its highly elastic nature and ease of molding. The PU masters were plasma-treated with O□plasma for 5 minutes, then functionalized with perfluorooctyl silane before Ecoflex molding. After the Ecoflex had cured for 90 minutes, the molds were peeled off. The same surface treatment was repeated for the Ecoflex negatives. The treated negatives were then used as molds to form the final PDMS positives. Each scaffold contained four wells with posts (diameter 0.4 mm × height 3.76 mm) spaced 4.68 mm apart. The posts of the scaffold were capped using 3D-printed ABS constructs to prevent slippage of engineered skeletal muscles (ESMs). The caps were secured with Ecoflex, autoclaved, and stored in sterile Milli-Q water. Representative images of the PU, Ecoflex, and PDMS stages and their dimensions are shown in Figure 1A.

**Figure 1.**
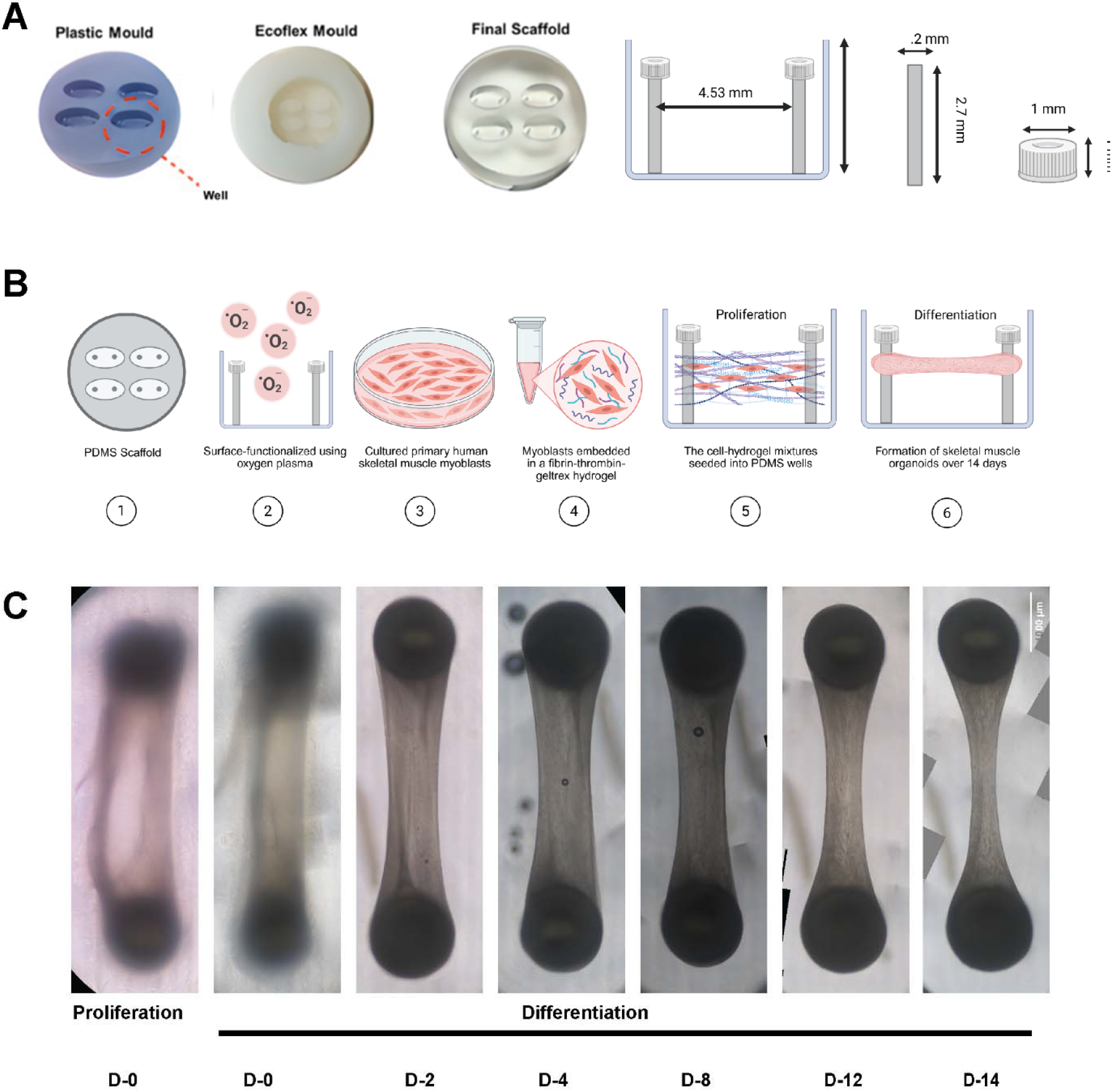
Design And Fabrication of a PDMS scaffold for use in Engineered Skeletal muscle culture. (A) Fabrication workflow: PU master used to cast an Ecoflex® negative, followed by the final polydimethylsiloxane (PDMS) scaffold bearing four wells; right, side-view schematic of a single well showing post geometry and well depth/width. Timeline for organoid generation from human myoblasts: surface functionalization, preparation of the cell–hydrogel mixture, seeding into the wells, culture and maturation (day P-1 to day 14, D14). Representative bright-field images of organoids at the indicated time points (P-1, D0, D2, D4, D8 and D12), illustrating progressive compaction and maturation. Scale bar, 1 mm.

### Generation of Human ESMs Using Primary Myoblasts in a Fibrin-Thrombin Hydrogel

Figure 1B shows the process of generating ESMs from human myoblasts embedded in a fibrin-thrombin hydrogel. Human skeletal muscle myoblasts were commercially acquired (Lonza, Cook MyoSite). Myoblasts from passage 6 to passage 8 were grown in proliferation medium (SKMGR). The cells were then trypsinized and mixed with one half of a two-part hydrogel. The hydrogel consisted of part A (cells, SKMGR, and thrombin) and part B (fibrinogen, SKMGR, and Geltrex). Components were kept separately on ice and combined immediately before seeding. Each ESM consisted of 400,000 myoblasts seeded into a single scaffold well and allowed to polymerize for one hour at 37°C. The ESMs were cultured in proliferation medium (SKMGR) for 4 days, then shifted to differentiation medium (SKMDR) for 14 days. Media was replenished every alternate day. Figure 1C shows the morphological changes from proliferation day 0 through differentiation days 0, 2, 4, 8, 12, and 14. ESMs progressively compacted, becoming narrower in appearance over the 14 days of differentiation.

### Immunohistological Characterization of Differentiated Skeletal Muscle Markers in Human ESMs

To assess the presence of differentiated skeletal muscle and myogenic markers in the fabricated ESMs, the following analyses were performed. ESMs were fixed with 4% paraformaldehyde overnight at 4°C. The fixed ESMs were then removed from the posts, immobilized in OCT medium, and sectioned at 10 µm resolution. Sections were stained with an anti-pan-MYHC antibody. The staining showed distinct myotubes distributed along the entire length of longitudinally sectioned ESMs (Fig. 2A). Nuclei stained with DAPI displayed elongated, aligned nuclei oriented along the same axis as the myotubes. The myotubes exhibited elongated, multinucleated syncytia with adjacent, distinct acetylcholine receptor (AChR) puncta.

**Figure 2.**
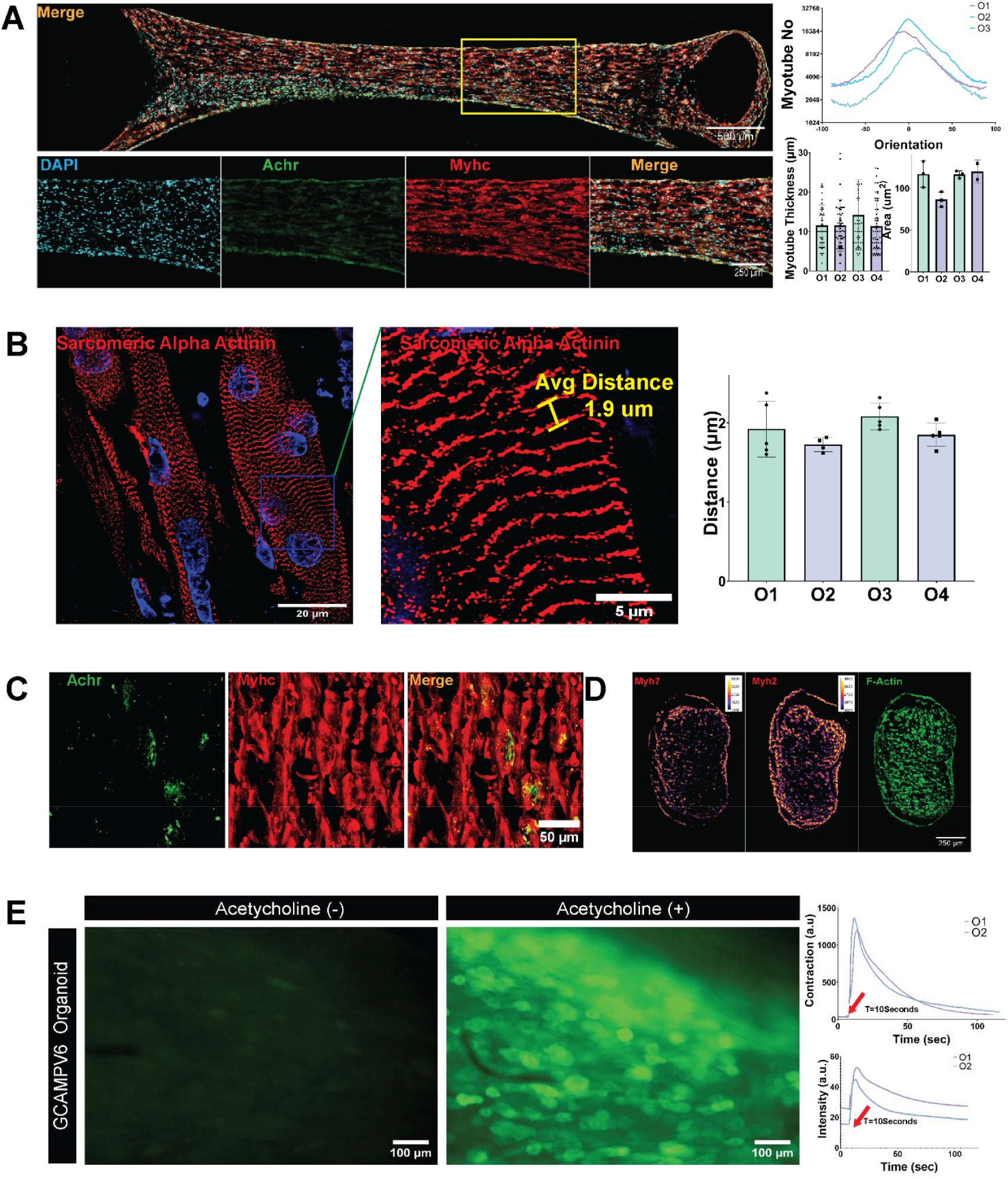
Immunohistochemical characterization of engineered skeletal muscle. (A) Longitudinal ESM section stained for nuclei (DAPI, cyan), acetylcholine receptors (AChR, green), and myosin heavy chain (MYHC, red); merged image shown. Boxed area is displayed below as single-channel images with merge. Right, bar graphs show the orientation of myotubes relative to the long axis, myotube thickness, and nuclear area across ESMs O1–O4. (B) Longitudinal ESM section stained for Sarcomeric Alpha Actinin, a higher-magnification region highlighting individual sarcomeres (average spacing ∼1.9 μm). Right, bar graph showing average intra-sarcomeric distance across O1–O4. (C) Longitudinal ESM section stained for AChR clusters (green) colocalized with MYHC-positive myotubes (red); merge at right. (D) Transverse ESM sections stained for MYHC isoforms—slow-twitch MYH7 and fast-twitch MYH2 and F-actin. (E) GCaMP6-expressing ESM before (ACh−) and after 250 μM acetylcholine treatment (ACh+); right, line graphs showing increased fluorescence intensity and contraction upon 250 μM acetylcholine treatment. Scale bars: (A, top) 500 μm; (A, bottom) 250 μm; (B, left) 20 μm; (B, right) 5 μm; (C) 50 μm; (D) 250 μm; (E) 100 μm.

To assess reproducibility across organoids, several metrics were analyzed from four ESMs and showed consistent morphology (Fig. 2A). The MYHC and DAPI signals were thresholded and segmented using ImageJ and analyzed using the MorphoLibJ and OrientationJ plugins. Orientation analysis demonstrated uniform myotube alignment along a single horizontal axis (n = 2 organoids). Myotube thickness showed no statistically significant variability across replicates (each data point represents one myotube; at least 50 myotubes were analyzed per ESM). Nuclear areas measured using the MorphoLibJ plugin showed no significant differences (each data point represents one field of view [FOV]; n = 3 FOVs per ESM). Quantitative shape analysis for convexity, perimeter, and circularity showed no significant differences among samples (each data point represents analysis from one FOV; three FOVs were analyzed per ESM to generate bar graphs in Fig. S2A).

### Formation of Sarcomeric Units, Adult Myosin Isoforms, and AChR Clusters

Longitudinal sections of ESMs were stained with an anti-sarcomeric α-actinin antibody (Abcam), which revealed distinctly formed sarcomeric units arranged in a repeating pattern (Fig. 2B). Average intra-sarcomeric distance was determined from line-profile analyses (10–20 sarcomeres per measurement; five measurements per organoid). Analysis revealed an average intra-sarcomeric distance of 1.9 µm, consistent with in vivo values, as shown in the adjacent bar graph.

Staining for AChRs revealed discrete clusters localized exclusively at the periphery of MYHC-positive myotubes (Fig. 2C). Transverse sections demonstrated the presence of adult myosin isoforms MYH2 (fast-twitch) and MYH7 (slow-twitch), which colocalized with myotubes visualized by F-actin staining (Fig. 2D). Transverse sections stained for pan-MYHC and laminin showed the presence of myotubes with well-defined laminin boundaries (Fig. S2B).

### Calcium Uptake in Response to Acetylcholine Treatment

To assess functional maturity, ESMs fabricated from GCaMP6-lentivirus-transduced myoblasts were used to measure calcium uptake (Fig. 2E). On day 14, ESMs from GCaMP-transduced myoblasts (n = 2) were treated with 250 µM acetylcholine. Upon treatment at t = 10 s, the ESMs exhibited an increase in fluorescence intensity, indicating calcium uptake. This fluorescence increase was accompanied by corresponding contraction. Video S1 displays both the increase in contraction and the corresponding fluorescence signal spike, confirming the presence of calcium-dependent contractile activity in the ESMs.

### NRH Increases NAD+ Levels in vitro and in vivo

The schematic in Figure S3A depicts two entry routes for dihydronicotinamide riboside (NRH): (1) a salvage route in which NRH is phosphorylated by an adenosine kinase-like “NRH kinase” to NMNH, then adenylated by NMNAT to NADH/NAD+; and (2) an oxidative route in which NRH is non-enzymatically oxidized to NR, phosphorylated by NRK to NMN, and converted by NMNAT to NAD+.

We confirmed that NRH increased intracellular NAD+ using multiple cell lines. HepG2 cells treated with NRH (5–250 µM, 4 h) exhibited concentration-dependent increases in intracellular NAD+ relative to untreated controls (Fig. S3B). HEK293 cells treated with NRH for 4 h (125, 250, and 500 µM) showed significantly higher NAD+ content than untreated cells (Fig. S3C). Jurkat cells treated for 4 h with either NRH or NR (5–1000 µM) showed a dose-dependent increase in NAD+ levels (Fig. S3D). NRH was significantly more effective in increasing intracellular NAD+ levels at this time point. We next determined the ability of NRH to increase NAD+ levels *in vivo*. Male Sprague-Dawley rats received intravenous NRH (50, 125, 250 mg/kg) at 0 h and 24 h; tissues were collected 4 h after the second dose. LC-MS/MS analysis showed higher NAD+ levels than vehicle in whole blood (Fig. S3E) and skeletal muscle (Fig. S3F) at 28 h after the first dose. Skeletal muscle NAD+ levels were significantly increased compared to vehicle (p = 0.035). Treatment with concentrations up to 500 µM did not show significant effects on cellular viability as judged by the MTT assay (Figure S3G). Based on these findings, we selected 500 µM as the test concentration for toxicology assays in human ESMs.

### Rationale for 500 µM NRH Concentration in 3D Organoid Studies

3D organoids present additional diffusion barriers and binding sinks compared with monolayers, so the nominal bath concentration overestimates the actual intratissue exposure, particularly in the core of the construct. Using 500 µM at the medium level compensates for these gradients and ensures that a substantial fraction of myofibers experience NRH levels comparable to those that maximally raise NAD+ in 2D systems. At the same time, this concentration should be viewed as a deliberately conservative, supra-physiological exposure—selected to define the upper boundary of our platform’s dynamic range and to map early structural and functional liabilities of extreme NAD+ boosting, rather than to mimic expected clinical plasma Cmax values, which remain incompletely characterized for NRH. In line with standard *in vitro* toxicology practice, using a high, non-lethal concentration such as 500 µM allows us to (1) confirm that the 3D skeletal muscle organoids can detect subcellular and functional toxicity endpoints, and (2) provide a benchmark against which future studies at lower, more clinically relevant exposures can be interpreted.

### Treatment with 500 µM NRH Increased Myotube Fusion and Total Nuclear Number

We assessed the effect of 500 µM NRH on skeletal muscle differentiation by measuring fusion index, a well-established metric of differentiation. 10-µm longitudinal ESM sections were stained for MYHC using an anti-MYHC pan-myosin antibody (MF-20, DSHB) alongside nuclei (DAPI) and acetylcholine receptor (AChR). Fusion index (Fig. 3A) was quantified using the MATLAB application ‘Myotube Analyser’. The software used both DAPI and MYHC channels and measured the number of nuclei colocalized with segmented MYHC. Fusion index is a percentage-based metric representing the proportion of nuclei within myotubes in a given field.

**Figure 3.**
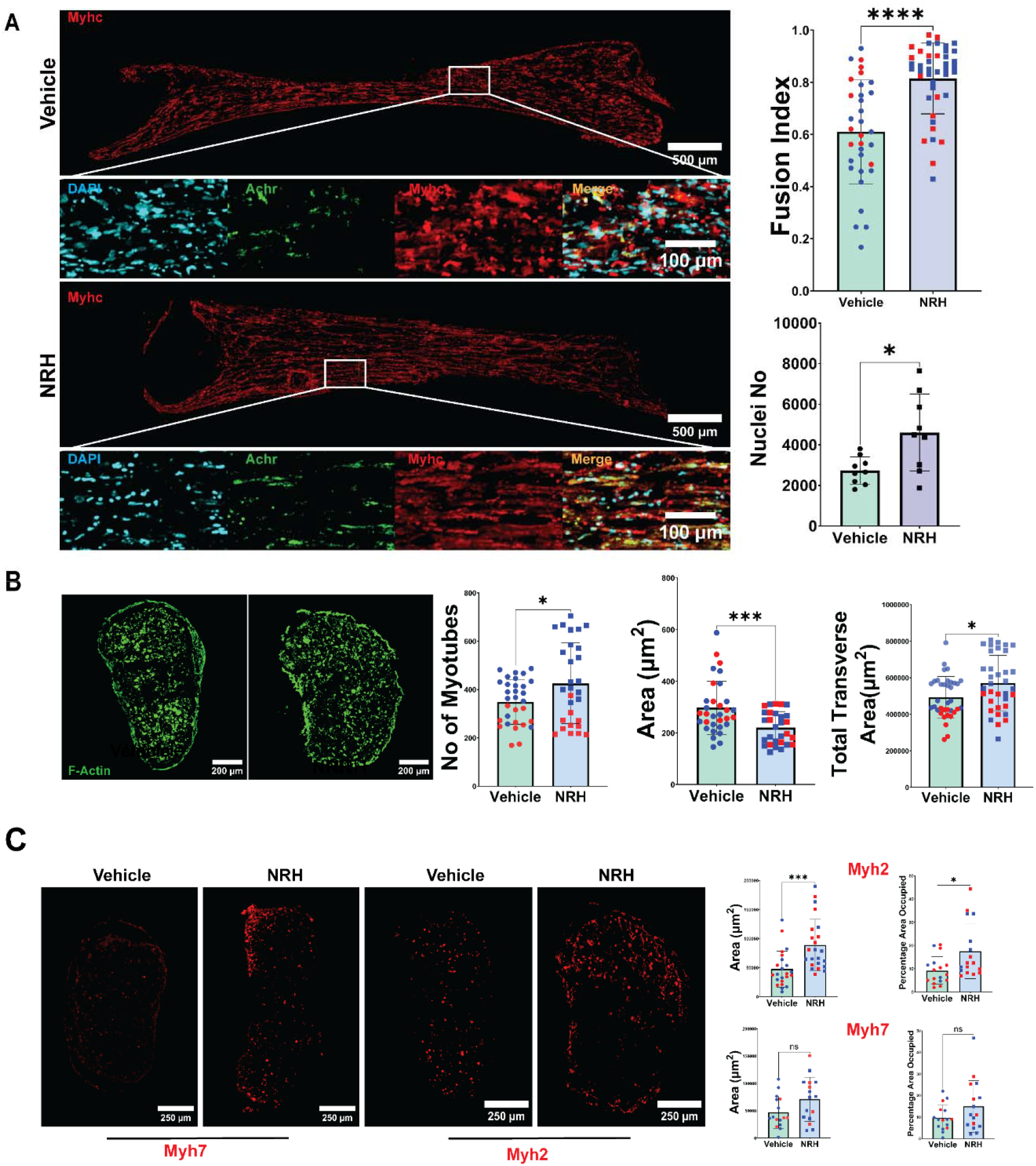
Vehicle versus NRH treatment in engineered skeletal muscles. (A) Longitudinal sections stained for DAPI (nuclei, cyan), AChR (green), and MYHC (red); Boxed area is displayed below as single-channel images with merge. Right, Bar graphs show Fusion index and nuclei per Longitudinal section (each point, one full section; points color-coded by donor: blue, donor 1 - 3 ESMs; red, donor 2 - 2 ESMs). Statistics (two-tailed, unpaired Student’s *t*-test): fusion index, **** *P* < 0.0001; nuclei per section, *P* < 0.05.(B) Transverse sections stained for F-actin. Right bar graphs of myotube number per section, mean myotube cross-sectional area, and total transverse area. Statistics (two-tailed, unpaired Student’s *t*-test): myotube number, *P* < 0.05; mean myotube area, *** *P* < 0.001; total transverse area, *P* < 0.05.(C) Transverse sections stained for MYH7 and MYH2. Right, bar graphs of isoform-positive area and percentage area compared to whole ESM occupied. Statistics (two-tailed, unpaired Student’s *t*-test): MYH2-positive area, *** *P* < 0.001; MYH2 % area, *P* < 0.05; MYH7-positive area, n.s.; MYH7 % area, *P* < 0.05. Scale bars: (A) 500 μm (top), 100 μm (bottom); (B) 200 μm; (C) 250 μm.

Quantitation of fusion index showed a statistically significant difference (p < 0.0001) between vehicle and 500 µM NRH-treated samples (each data point represents one FOV; minimum of 30 FOVs were analyzed per condition). A minimum of 6 FOVs were chosen per ESM to sample the entirety of the structure. This ensures that any gradient of efficacy could be captured if present and would not present a confounding variable for the analysis. ESMs were engineered using myoblasts derived from two donors. Data points are color-coded: blue for donor 1, red for donor 2. The DAPI channel was segmented and counted using the MorphoLibJ plugin and was found to be significantly elevated in the NRH treatment group (Fig. 3A, adjacent bar graph; p < 0.05; each data point represents one FOV; minimum of 3 FOVs per ESM were analyzed).

### NRH Treatment Alters ESM Morphology: Increased Total Area and Myotube Number, Decreased Individual Myotube Size

To evaluate NRH effects on myotube morphology (Fig. 3B), we labelled 10-µm transverse ESM sections for filamentous actin (F-actin) and imaged under identical settings across treatment groups. Segmentation of F-actin-positive myotubes was performed to compute three specific metrics per section: (1) number of myotubes, (2) mean myotube cross-sectional area (µm^2^), and (3) total transverse area (µm^2^) occupied by ESM myotubes. NRH significantly increased both the number of myotubes and the total transverse area of the ESM but reduced individual myotube size (Student’s t-test; p < 0.05 for myotube number, p < 0.001 for mean area, p < 0.05 for total transverse area). Each data point represents one FOV; a minimum of 30 FOVs per condition were analyzed. Data points are color-coded: blue for donor 1, red for donor 2.

### Treatment with 500 µM NRH Selectively Increased MYH2-Positive Myotube Area and Coverage

We next assessed fibre-type composition on 10-µm transverse sections using isoform-specific myosin antibodies (Fig. 3C): fast-twitch type IIa myosin heavy chain (MYH2; SC-71, DSHB) and slow-twitch type I myosin heavy chain (MYH7; A4.95, DSHB). For each section, we quantified (1) isoform-positive area (µm^2^) and (2) percent area coverage within the construct.

NRH significantly increased MYH2-positive area and coverage relative to vehicle (Student’s t-test; p < 0.001 for area, p < 0.05 for percent coverage), whereas MYH7-positive area and coverage showed no significant change (p > 0.05). Each data point represents one FOV; minimum of 10 FOVs per condition. Data points are color-coded: blue for donor 1, red for donor 2.

### NRH Treatment Increased Mitochondrial Area Without Significantly Altering Membrane Potential

To analyze the effects of NRH on mitochondrial structure and function, we conducted live-cell mitochondrial imaging of ESMs. Whole, live day-16 ESMs were stained with tetramethylrhodamine methyl ester (TMRM) and imaged using live-cell confocal microscopy. This probe accumulates in polarized mitochondria, allowing quantification of mitochondrial membrane potential changes.

The TMRM signal was segmented using ImageJ and a mask was created. The mask was used to measure mitochondrial intensity only for the stained area (signifying mitochondrial potential) for vehicle versus NRH 500 µM-treated samples. Bar graph (Fig. 4A) shows a non-significant trend toward increased mitochondrial potential (Student’s t-test, p > 0.05). The segmented mask was also used to quantify the area of individual mitochondria. Figure 4A (left) shows a line graph displaying the distribution of mitochondrial areas (n = 3 ESMs per condition): blue line represents vehicle, red line represents NRH-treated ESMs. We observed an increase in the average area of mitochondria when treated with 500 µM NRH, suggesting increased mitochondrial size upon NRH treatment.

**Figure 4.**
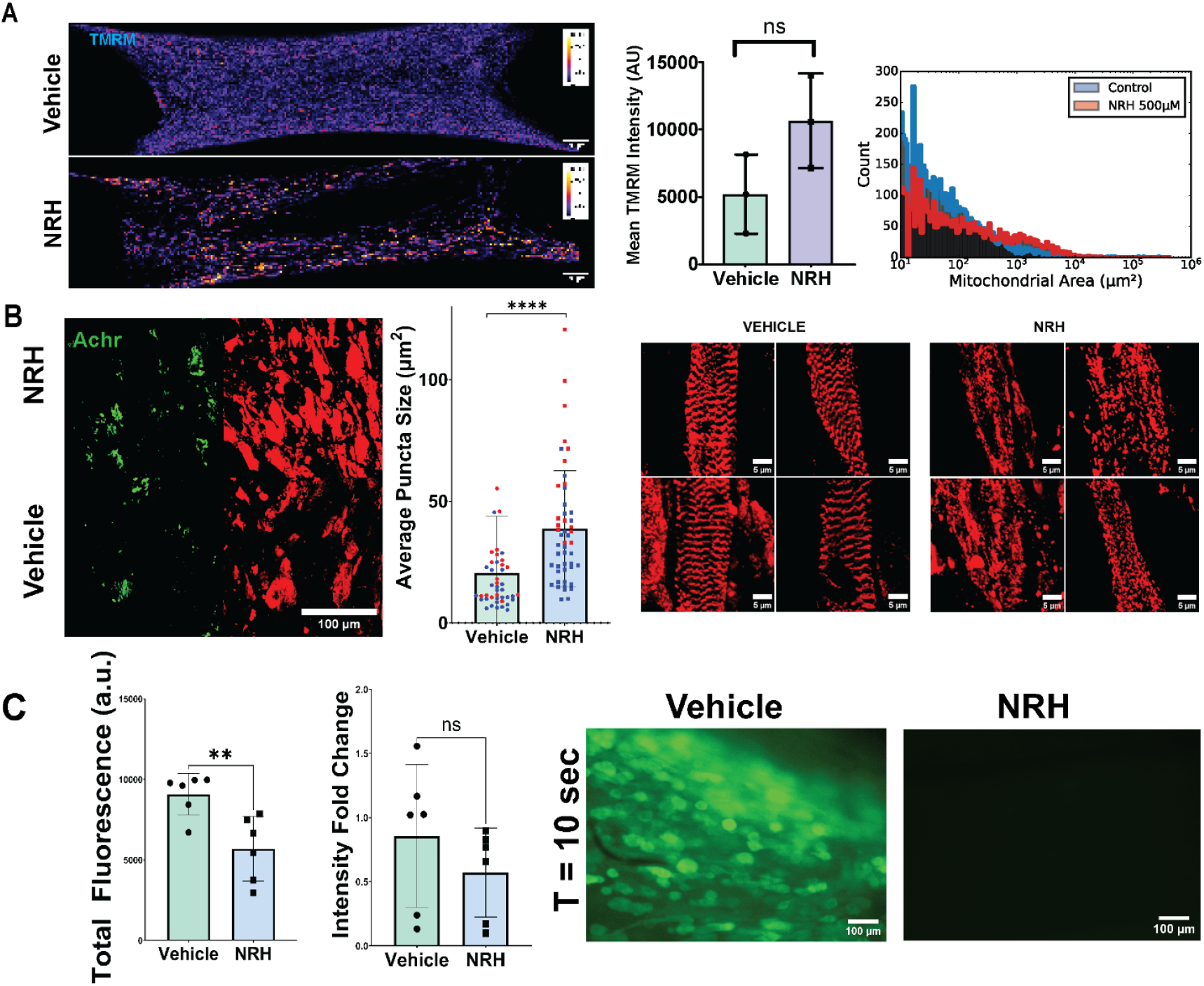
Functional effects of NRH (500 μM) in ESMs. (A) Whole ESMS stained with TMRM (pseudocolor intensity map). Right, bar graph of mean TMRM intensity per ESM and Frequency distribution of mitochondrial area (Each line represents 2 ESMs). Statistics (two-tailed, unpaired Student’s *t*-test): mean TMRM intensity, n.s.(B) Longitudinal sections stained for AChR (green) and MYHC (red). Centre, bar graph of average AChR puncta size per section (each point, one full section; points color-coded by donor: blue, donor 1 - 3 ESMs; red, donor 2 - 2 ESMs). Statistics (two-tailed, unpaired Student’s *t*-test): average puncta size, **** *P* < 0.0001. Right, Longitudinal ESM sections stained for Sarcomeric Alpha Actinin in vehicle and NRH Treatment (donor 1 - 2 ESMs; red, donor 2 - 2 ESMs). (C) Bar graphs summarizing GCaMP6 calcium responses to acetylcholine (250 μM): left, total fluorescence (area under the curve) per ESM; centre, fold-change in fluorescence intensity relative to baseline (each point, one ESM). Statistics (two-tailed, unpaired Student’s t-test): total fluorescence, **P < 0.01; intensity fold change, n.s. right, snapshot at *T* = 10 s from representative vehicle and NRH treated ESMs. Scale bars: (A) 250 μm; (B, left) 100 μm; (B, right) 5 μm; (C) 100 μm.

### Treatment with 500 µM NRH Increased AChR Puncta Size and Disrupted Sarcomeric Organization

10-µm longitudinal ESM sections were co-stained for acetylcholine receptor (AChR; α-bungarotoxin Alexa Fluor 488) and MYHC. AChR puncta were segmented using FIJI and their areas quantified. NRH treatment over 14 days significantly increased the average AChR puncta size relative to vehicle (Student’s t-test, p< 0.0001; each data point represents one section; n ≥ 30 sections per condition; data points color-coded by donor as described above), indicating enlarged receptor clusters. Longitudinal sections were stained with sarcomeric α-actinin (Abcam), and high-magnification confocal imaging revealed clearly formed sarcomeric banding in controls, whereas NRH-treated samples exhibited disrupted sarcomere assembly (Fig. 4B).

### Treatment with 500 µM NRH Inhibited Acetylcholine-Induced Calcium Signaling

We quantified calcium signaling dynamics in ESM constructs stably expressing a genetically encoded calcium indicator (GCaMP) delivered by lentiviral transduction. GCaMP fluorescence was recorded over time following treatment of 250 µM acetylcholine at t = 10 s from a defined region of interest. NRH-treated constructs showed a significant reduction in total GCaMP fluorescence (time-integrated signal) compared to vehicle controls, while the peak fold-change in fluorescence relative to baseline was not significantly different between groups (Fig. 4C; each point represents one ESM; two-tailed unpaired Student’s t-test, total fluorescence **P < 0.01, fold-change n.s; imaging settings were identical across groups). Representative imaging showed robust calcium activation in vehicle-treated ESMs and minimal detectable signal in NRH-treated constructs (Fig. 4C).

## Discussion

This study establishes 3D Engineered human skeletal muscle organoids as a physiologically relevant platform to evaluate the toxicological effects of NAD+-boosting compounds such as dihydronicotinamide riboside (NRH). At a deliberately high concentration (500 µM) over 14 days, NRH produced a dual phenotype: it enhanced classical markers of myogenic differentiation yet simultaneously induced significant structural and functional defects. NRH increased fusion index, total myotube number, and selectively expanded fast-twitch (MYH2-positive) fiber area, but it also reduced individual myotube size, disrupted sarcomeric organization, enlarged acetylcholine receptor (AChR) puncta, and markedly blunted acetylcholine-evoked calcium transients. These findings highlight the value of 3D organoids for uncovering complex, dose-dependent effects of NRH that may be missed in conventional 2D systems.

### NRH-enhanced differentiation with impaired maturation

NRH treatment increased both the fusion index and nuclear number, indicating that it accelerates myoblast fusion and commitment to the myogenic lineage. The fusion index, which quantifies the proportion of nuclei within multinucleated myotubes, is a readout of myogenic differentiation. The observed increase in fusion index suggests that elevated NAD+ may promote progression through early myogenesis. NAD+-dependent regulators such as sirtuins and myogenic transcription factors are known to play a roles in skeletal muscle homeostasis (Fulco, Schiltz et al. 2003, Goody and Henry 2018). This apparent benefit was counterbalanced by a trade-off. NRH increased the number of myotubes and total cross-sectional area but reduced individual myotube size. Rather than generating larger, well-organized fibers, NRH produced more numerous, smaller myotubes with compromised sarcomeric organization. Muscle force is driven by fiber cross-sectional area and sarcomere integrity. These changes may impact functional properties. Our data emphasize that reliance on fusion index or myotube number alone can be misleading. Multidimensional readouts that capture both structural integrity and function are essential for evaluating myogenic interventions.

### Fiber-type specificity and metabolic implications

NRH selectively increased the area and coverage of MYH2-positive (fast-twitch type IIa) fibers without significantly altering MYH7-positive (slow-twitch type I) fibers. This fiber-type bias suggests that NRH modulates myogenic or metabolic programs in a fiber-specific manner. Fast-twitch fibers are more glycolytic and less oxidative than slow-twitch fibers. A preferential expansion of MYH2-positive fibers may reflect changes in fiber-type specification associated with NRH treatment.

Clinically, preservation of fast-twitch fibers may be beneficial in aging. Type II fiber loss contributes to reduced power and increased fall risk (Deschenes 2004, Kramer, Snijders et al. 2017). However, the potential advantage of selectively expanding MYH2 fibers depends on their structural and functional integrity. Since concurrent sarcomeric disorganization and calcium signaling defects we observed, the NRH-induced MYH2 expansion may may reflect fibers with altered structural and functional properties.

### Structural and functional defects: sarcomeres, AChR clusters, and calcium signaling

The key phenotypes in NRH-treated organoids were the disruption of sarcomeric organization and the marked reduction in acetylcholine-induced calcium signaling. In control organoids, α-actinin staining revealed well-organized sarcomeres with physiological spacing (∼1.9 µm), whereas NRH-treated constructs displayed fragmented and disordered sarcomeric patterns. Because sarcomeres are the fundamental contractile units of muscle, their disruption signals a major functional liability that cannot be compensated by increased myotube number.

Functionally, NRH-treated ESMs exhibited a reduction in acetylcholine-evoked calcium transients despite significantly enlarged AChR puncta. This uncoupling between receptor clustering and downstream calcium signaling could suggest defects at one or more nodes of the excitation–contraction (EC) coupling cascade. This could include AChR function, voltage-gated calcium channels, ryanodine receptors, or calcium reuptake and buffering mechanisms. The enlargement of AChR clusters could be associated with impaired signaling.

Mitochondrial analyses revealed increased mitochondrial area with only a non-significant trend toward elevated membrane potential. These data suggest morphological remodelling without clear functional gain. This could also reflect stress responses such as swelling or incomplete biogenesis. Because sarcomere assembly and calcium handling are energetically demanding, changes in mitochondrial structure may be relevant to the observed structural and functional phenotypes.

### Mechanistic integration: the NAD+ –redox–calcium axis

Our findings are consistent with the possibility that supra-physiological NRH remodels skeletal muscle biology through effects on NAD+ metabolism, redox balance, and calcium-dependent signaling:

i. Excessive NAD+ elevation and sirtuin/PARP dysregulation-Very high NAD+ levels can hyperactivate sirtuins and alter deacetylation of cytoskeletal, metabolic, and transcriptional regulators (Houtkooper, Pirinen et al. 2012, Poljsak, Kovac et al. 2022), while PARP activity can reshape gene expression and consume NAD+ (Ryu, Kim et al. 2015, Murata, Kong et al. 2019, Hurtado-Bages, Knobloch et al. 2020). Together, these processes may disturb programs controlling differentiation, sarcomere assembly, and calcium handling.
ii. Redox perturbation and metabolic reprogramming-NRH enters the cell in reduced form and can elevate NADH as well as NAD+, potentially disturbing redox balance, electron transport chain function, and ROS production. These effects may drive a glycolytic shift and contribute to the selective expansion of MYH2 fibers while impairing oxidative capacity and ATP supply required for maturation.
iii. Calcium signaling disruption and feedback on structure-The strong reduction in acetylcholine-evoked calcium signals implicates broad disruption of calcium-dependent pathways that regulate fiber-type specification, sarcomere assembly and AChR clustering.

These mechanisms are not mutually exclusive and collectively highlight a window between beneficial and deleterious NAD+ elevation in muscle.

### Advantages of 3D skeletal muscle organoids for toxicology

A key question is why these NRH-induced defects have not been widely reported. We propose that the 3D organoid context was critical for their detection. First, mechanical loading between posts in our scaffold imposes stringent requirements for sarcomere alignment and force transmission that are absent in 2D monolayers, making structural defects more apparent. Second, simultaneous assessment of calcium dynamics and contraction in mechanically constrained tissue provides a more faithful readout of EC coupling than 2D cultures. Third, diffusion gradients within 3D constructs generate heterogeneous exposure, approximating *in vivo* conditions more closely than uniform 2D systems and revealing concentration-dependent effects across a range of local NRH levels. Finally, the extended differentiation period only possible in 3D cultures, and multidimensional endpoints (morphology, fiber type, mitochondria, calcium) allowed us to distinguish early differentiation gains from later maturation failures.

Collectively, these features position ESMs as an intermediate-complexity system between 2D cultures and animal models, with enough physiological relevance to uncover subtle but functionally important toxicities.

### Limitations and implications for NAD+ precursor development

We tested a single, high NRH concentration (500 µM) chosen to define an upper bound of toxicity and to overcome diffusion limitations. Clinically relevant exposures are likely lower and remain incompletely characterized. ESMs were derived from only two donors, therefore inter-individual variability in response to NRH is not yet defined. Our analyses focused on a single time point (day 14) and did not include longitudinal or reversibility studies, nor comprehensive multi-omics profiling. In addition, the organoids lack vasculature, innervation, and immune components, and therefore cannot capture systemic pharmacokinetics, inflammatory responses, or neuromuscular interactions. Despite these constraints, the data show a proof-of-concept that high-level NAD+ boosting by NRH can uncouple early differentiation from proper structural and functional maturation in 3D human skeletal muscle organoids.

In conclusion, our data indicate that 500 µM NRH enhances early myogenic differentiation and induces structural and functional defects in 3D human skeletal muscle organoids. This dual phenotype illustrates the complexity of targeting NAD+ metabolism in muscle, this shows the limitations of relying on NAD+ levels or simple differentiation metrics as sole indicators. By analyzing fiber-type-specific effects, sarcomere disruption, changes in mitochondria, and impaired calcium uptake, this work suggests that 3D skeletal muscle organoids may be useful for assessing the effects of metabolic interventions such as NRH.

## Data availability statement

The original contributions presented in the study are included in the article/supplementary material, further inquiries can be directed to the corresponding author.

## Author contributions

AR: Validation, Methodology, Data curation, Visualization, Supervision, Project administration, Conceptualization, Investigation, Formal Analysis, Writing – original draft, Writing – review and editing, AV-Validation, Methodology, Data curation, Visualization, Investigation, Formal Analysis, Writing – original draft, Software, SSP – Investigation, YS: Investigation, AH: Investigation, BV-Methodology, SSD: Investigation, AB: Investigation, HL: Methodology, SCL: Investigation, Methodology, NZ: Data curation, GP: Formal Analysis, GMS: Conceptualization, Formal Analysis, Writing – review and editing.

## Funding and Ethics statement

The study was funded by grant from Mitosciences LLC. and Department of Biotechnology, Govt. of India. All ethical guidelines were followed as required by the journal.

## Conflict of interest

Three coauthors affiliated with MitoPower Inc., a company that is investigating NRH for potential use in human disease. Their involvement does not alter the authors’ adherence to all journal policies on data sharing, analysis, and reporting. All experimental work, data interpretation, and manuscript preparation were carried out without commercial influence, and the company had no role in study design, data collection, or decisions regarding publication. The remaining authors declare no competing financial interests.

## Generative AI statement

The authors declare that the Nature Research Assistant AI tool was used to assist in and editing the manuscript

## Notes

### Competing Interest Statement

The authors have declared no competing interest.

