## Supplementary Figures for "3D human skeletal muscle organoids reveal distinct effects of high-dose dihydronicotinamide riboside on muscle development"

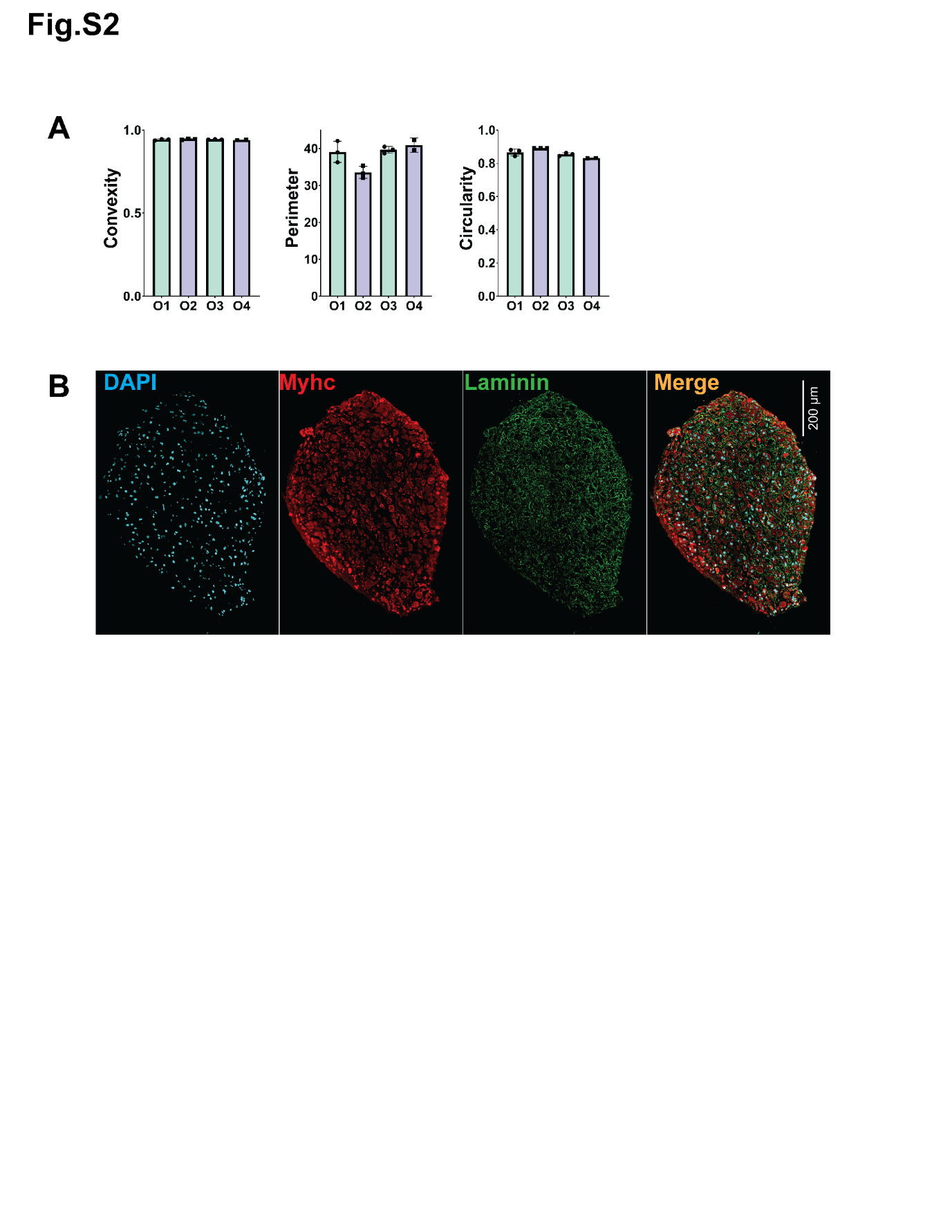

**Supplementary Figure S2. Nuclear shape analysis and transverse section morphology of engineered skeletal muscles.** (A) Bar graphs of Nuclear Shape—convexity, perimeter, and circularity—measured across organoids O1–O4 (each point, one full section). Statistics: one-way ANOVA. (B) Representative transverse section stained for DAPI (nuclei, cyan), MYHC (red), and laminin (green), merge at right. Scale bar, 200 μm.

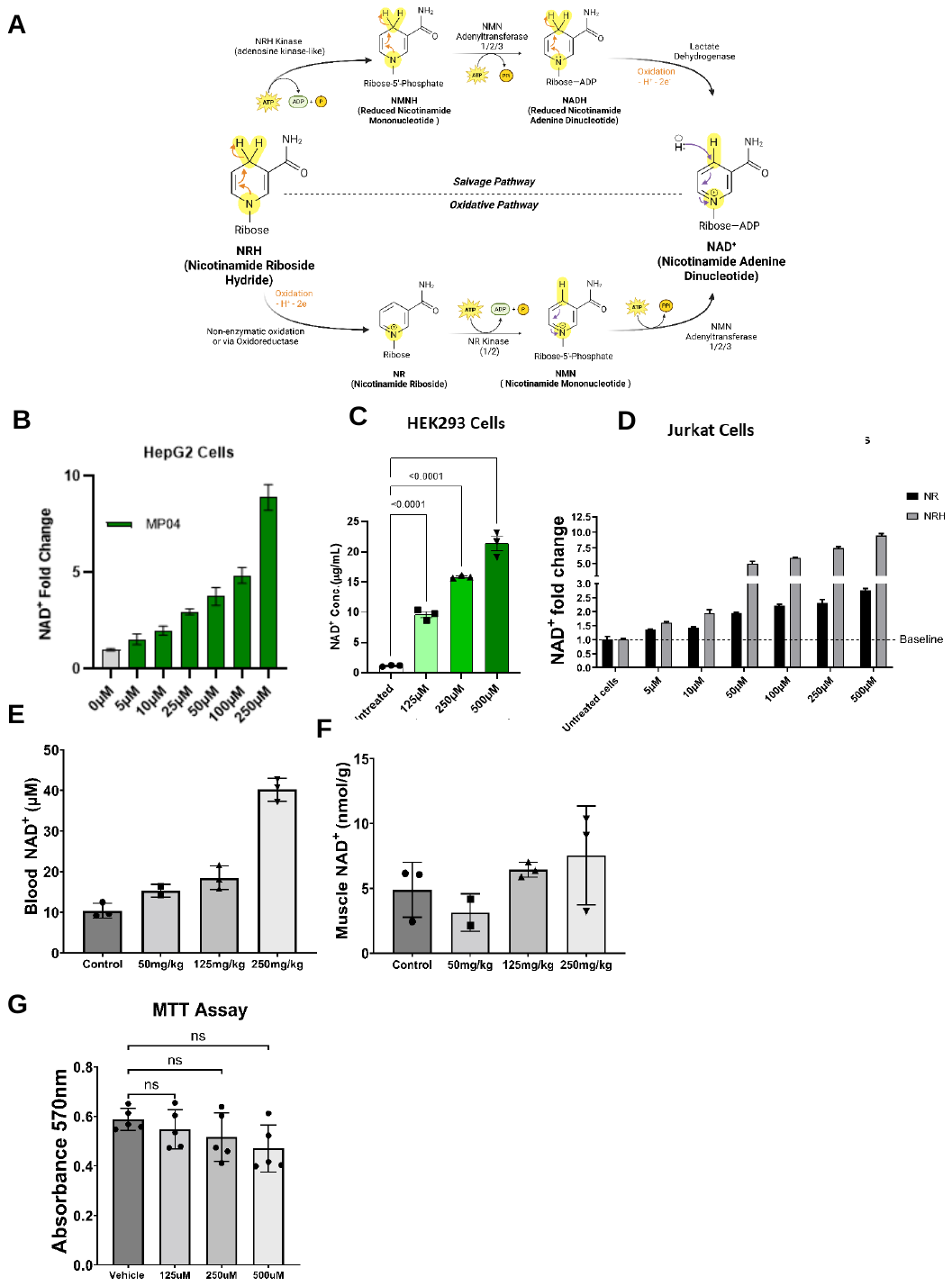

**Supplementary Figure S3. NRH increases cellular and tissue levels of NAD⁺.** (A) Schematic of the NRH pathway overview. (B,C.D) Bar graphs of intracellular NAD⁺ levels measured by LC–MS/MS in human HepG2, HEK293, and Jurkat cells after treatment with NRH (5–500 μM, 4 h) or NR (5–1000 μM, 4 h).(E,F) Bar graphs of tissue NAD⁺ levels in male Sprague–Dawley rats following two doses of NRH (50–250 mg kg⁻¹, 0 h and 24 h), with terminal collection at 4 h after second dose skeletal muscle (nmol g⁻¹), and whole blood (μM). (G) Bar graph of C2C12 myoblasts MTT assay (absorbance 570 nm) following NRH (125–500 μM) treatment
